# Machine learning uncovers a data-driven transcriptional regulatory network for the Crenarchaeal thermoacidophile *Sulfolobus acidocaldarius*

**DOI:** 10.1101/2021.07.28.454237

**Authors:** Siddharth M. Chauhan, Saugat Poudel, Kevin Rychel, Cameron Lamoureux, Reo Yoo, Tahani Al Bulushi, Yuan Yuan, Bernhard O. Palsson, Anand V. Sastry

## Abstract

Dynamic cellular responses to environmental constraints are coordinated by the transcriptional regulatory network (TRN), which modulates gene expression. This network controls most fundamental cellular responses, including metabolism, motility, and stress responses. Here, we apply independent component analysis, an unsupervised machine learning approach, to 95 high-quality *Sulfolobus acidocaldarius* RNA-seq datasets and extract 45 independently modulated gene sets, or iModulons. Together, these iModulons contain 755 genes (32% of the genes identified on the genome) and explain over 70% of the variance in the expression compendium. We show that 5 modules represent the effects of known transcriptional regulators, and hypothesize that most of the remaining modules represent the effects of uncharacterized regulators. Further analysis of these gene sets results in: (1) the prediction of a DNA export system composed of 5 uncharacterized genes, (2) expansion of the LysM regulon, and (3) evidence for an as-yet-undiscovered global regulon. Our approach allows for a mechanistic, systems-level elucidation of an extremophile’s responses to biological perturbations, which could inform research on gene-regulator interactions and facilitate regulator discovery in *S. acidocaldarius*. We also provide the first global TRN for *S. acidocaldarius*. Collectively, these results provide a roadmap towards regulatory network discovery in archaea.

## Introduction

Over the past few decades, scientists have been increasingly intrigued by the remarkable microorganisms that reside in extreme environments. The phenotypic diversity of these extremophiles is high, and due to the environments they inhabit, challenging to study. However, in recent years, advances in various molecular biology approaches – low-cost sequencing in particular – have enabled the exploration of the genotypic space these organisms inhabit.

*Sulfolobus acidocaldarius* is one such extremophilic archaeon; it is a thermoacidophile that resides in sulfur hot springs, which have an average pH of 2.4 and an average temperature of 83°C (Brock et al., 1972; Lewis et al., 2021). Such extremes in acidity and temperature necessitate specific adaptations to thrive in these habitats. *Sulfolobus* species are known to have an extraordinarily malleable yet chemiosmotic-resistant cell envelope, as well as exceptionally thermostable enzymes (Albers and Meyer, 2011). These characteristics make *Sulfolobus* valuable potential sources of biologics for future medical and biotechnological applications (e.g. production of carbocyclic nucleotides, halogen group removal, esterification) (Littlechild, 2015; Quehenberger et al., 2017). *S. acidocaldarius* is also one of the best-studied archaea as it is one of the few genetically tractable archaeal model systems (Wagner et al., 2012). While much progress has been made using these approaches, especially with regards to RNA-seq studies that have found multiple regulators of gene expression (Song et al., 2013-3; Reimann et al., 2012; Lassak et al., 2013; Liu et al., 2016; Haurat et al., 2017; Lemmens et al., 2019; Wang et al., 2019), much of the TRN structure of *S. acidocaldarius* remains unknown. Here, we leverage machine learning to construct a global TRN for this extremophile.

Independent Component Analysis (ICA) (Jutten and Herault, 1991) is an unsupervised machine learning algorithm that is especially useful for blind source separation of mixed signals. This algorithm takes in observations of mixed signals and is able to deconvolute them back into their independent, unmixed forms with no additional inputs (Hyvärinen and Oja, 2000). This methodology has proven quite successful in deconvoluting observed gene expression profiles into linear combinations of statistically independent genetic modules as the blind signals (Teschendorff et al., 2007; Biton et al., 2014; Sastry et al., 2019; Sompairac et al., 2019; Poudel et al., 2020; Rychel et al., 2020). These independently modulated sets of genes (termed iModulons) contain anywhere from one to hundreds of genes, and are often controlled by a common regulatory source, such as a single transcription factor (TF) or a combination of regulatory elements acting in concert (Sastry et al., 2019). iModulons containing only one gene often form as a result of noise or genetic perturbation in the compendium (e.g. gene knockout) (Sastry et al., 2021a).

In contrast to regulons, which are bottom-up groupings of co-regulated genes derived from transcription factor-to-DNA binding assays and other biomolecular methods, iModulons are co-expressed sets of genes which are derived from applying data analytics to large transcriptomic compendia. Thus, iModulons can be interpreted as data-driven analogs of regulons. Furthermore, the condition-dependent activity levels of each iModulon correspond to the activity of the underlying genetic signals and regulators. Despite the difference in approach (i.e. data analytics vs direct molecular methods), iModulons recapitulate many known regulons from the literature, and have accurately predicted new genetic targets for regulators (Sastry et al., 2019; Poudel et al., 2020; Rychel et al., 2020) and elucidated gene functions (Rodionova et al., 2020, 2021). This top-down approach allows for an unbiased method for reconstructing the TRN of prokaryotic organisms of interest, including archaea. As iModulons enable us to learn the TRN structure from transcriptomes alone, they represent a valuable approach for quickly characterizing relatively under-studied TRNs, such as that of *S. acidocaldarius*.

Given the considerable insights provided by ICA, along with a proven track-record (Sastry et al., 2019; Poudel et al., 2020; Rychel et al., 2020), we applied it to a compendium of 95 publically available RNA-seq expression profiles for *S. acidocaldarius* to deconvolute its TRN. This is the first application of ICA towards deconvolution of an archaeal TRN, and has led towards the generation of the most complete, global TRN of *S. acidocaldarius*. Our deconvolution reveals 45 robust iModulons, which explain 72% of the variance in the RNA-seq compendium. Of these, 5 iModulons strongly resemble well-characterized regulons with identifiable regulators. This top-down, systems-level view of the TRN allows us to use our iModulons to propose informed hypotheses, backed with computational evidence of our claims. We also present various groupings of poorly characterized genes which warrant further study, given their importance in regulation of gene expression. Static graphical summaries of all *S. acidocaldarius* iModulons, including enriched gene members and iModulon activities, are presented in the Supplementary Information, with interactive summaries available on iModulonDB.org (Rychel et al., 2021).

## Materials and Methods

Here, we demonstrate how we built the iModulon structure of *Sulfolobus acidocaldarius* from publicly available RNA-seq datasets (**Figure 1A**). The workflow followed here is based on the PyModulon workflow described by Sastry et. al. 2021 (Sastry et al., 2021b). All code to reproduce this pipeline is available at: https://github.com/SBRG/modulome_saci and https://github.com/avsastry/modulome-workflow/. Since this process results in the totality of iModulons that can currently be computed for this organism, we have named the resulting database as the “*S. acidocaldarius* Modulome’’.

**Figure 1:**
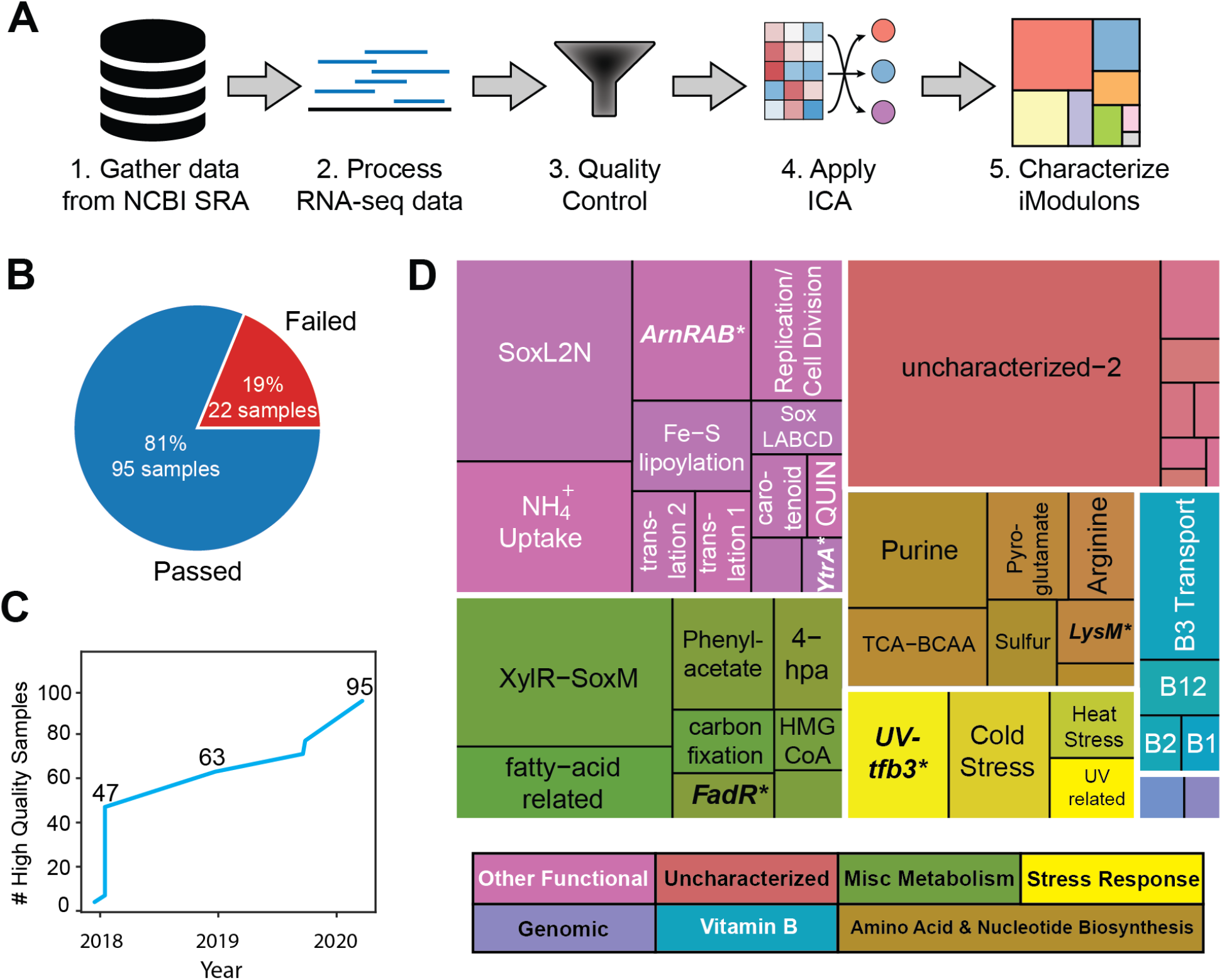
Overview of compiled *S. acidocaldarius* RNA-seq compendium and its iModulon characterization. **A**.) Graphical representation of the PyModulon ICA workflow (adapted from Sastry et al. (Sastry et al., 2021b)). **B**.) Number of samples that passed quality control. **C**.) Number of high-quality RNA-seq expression profiles for *S. acidocaldarius* in NCBI SRA over time. **D**.) Treemap of iModulons generated from the curated RNA-seq compendium. Sizes are representative of variance explained by iModulon (total 72%). Boldface and asterisk indicate that the corresponding iModulon recapitulates a known regulon.

### Gathering and Processing RNA-seq Data from NCBI SRA

Following the PyModulon workflow, we used a script that compiles the metadata for all publicly available RNA-seq data for a given organism in NCBI SRA (https://github.com/SBRG/modulome_saci/tree/master/notebooks/0_metadata). As of August 2020, we identified 117 expression profiles labelled as *Sulfolobus acidocaldarius* RNA-seq data. The FASTQ files for these datasets were collected and processed as previously described (Sastry et al., 2021b).

Briefly, we downloaded raw FASTQ files using fasterq-dump (https://github.com/ncbi/sra-tools/wiki/HowTo:-fasterq-dump), and performed read trimming using Trim Galore (https://www.bioinformatics.babraham.ac.uk/projects/trim_galore/) with the default options, followed by FastQC (http://www.bioinformatics.babraham.ac.uk/projects/fastqc/) on the trimmed reads. Next, reads were aligned to the genome using Bowtie(Langmead et al., 2009). The read direction was inferred using RSeQC (Wang et al., 2012) before generating read counts using featureCounts (Liao et al., 2014). Finally, all quality control metrics were compiled using MultiQC (Ewels et al., 2016) and the final expression compendium was reported in units of log-transformed Transcripts per Million (log-TPM).

To process the complete *S. acidocaldarius* RNA-seq compendium, we used Amazon Web Services (AWS) Batch to run the Nextflow pipeline (Di Tommaso et al., 2017) (https://github.com/avsastry/modulome-workflow/tree/main/2_process_data)

### Quality Control

A Jupyter notebook (IOS Press Ebooks - Jupyter Notebooks – a publishing format for reproducible computational workflows) showcasing the quality control workflow can be found at https://github.com/SBRG/modulome_saci/blob/master/notebooks/1_expression_QC_SOP.ipynb. The final high-quality *S. acidocaldarius* compendium contained 95 RNA-seq datasets (**Figure 1B, 1C**). As part of the quality control procedure previously described (Sastry et al., 2021b), we performed manual curation of experimental metadata to identify which samples were biological replicates. We also examined the literature to identify each sample’s strain, media, additional treatments, environmental parameters/changes, and growth stage, if reported. During curation, we removed some non-traditional RNA-seq datasets, such as non-coding RNA-seq.

A draft TRN was also constructed at this stage. This process involved curating all reported regulatory interactions from *S. acidocaldarius* literature into a standardized table for further analysis downstream. Literature used in metadata annotation also served as material for developing a draft TRN (**Supplementary File 1**).

To obviate any batch effects resulting from collating different expression profile datasets, we selected a reference condition (consisting of at least two replicates) to normalize each project in the compendium. This ensured that nearly all independent components generated were due to biological variation rather than technical variation. After normalization, however, gene expression and iModulon activities can only be compared within a project to a reference condition, rather than across projects.

### Gene Naming and Annotation

The genome annotation derived from NCBI (accession number: NC_007181.1) contains many poorly characterized gene products, so the genome fasta was reannotated using the software tool prokka (Seemann, 2014). This resulted in prokka-specific locus tags and prokka-specific annotations. All of this data was then collated into a standardized gene annotation table using the following Jupyter notebooks: https://github.com/SBRG/modulome_saci/blob/master/notebooks/3_a_gene_annotation-manual-curation-notebook.ipynb and https://github.com/SBRG/modulome_saci/blob/master/notebooks/3_b_gene_annotation.ipynb. We then manually parsed through the literature to find as many relevant gene names and functional annotations as possible, generating a curated gene annotation table for *S. acidocaldarius* (**Supplementary File 2)**.

### ICA

Following the PyModulon workflow, we implemented ICA using the *optICA* extension of the popular algorithm FastICA. The *optICA* script used can be found at https://github.com/avsastry/modulome-workflow/tree/main/4_optICA and produces two matrices. One matrix (the **M** matrix) contains the robust independent components (McConn et al., 2021), and the other (the **A** matrix) contains the corresponding activities. The product of the **M** and **A** matrices approximates the expression matrix (the **X** matrix), which is the curated RNA-seq compendium. Each independent component in the **M** matrix is filtered to find the genes with the largest absolute weightings, which ultimately generates iModulon gene sets.

Implementing this process resulted in 45 iModulons for the *S. acidocaldarius* Modulome that explained 72% of the expression variance in the compendium (**Figure 1D**). Although the fraction of variance explained by 45 iModulons is much lower than the fraction of variance explained by 45 principal components of the **X** matrix, explained variance has additional meaning in ICA compared to PCA. Principal components are a mathematical representation of the compendium and often lack biological interpretation. However, independent components (and the iModulons which derive from them) can be directly linked to transcriptional regulation (via characterization), revealing interpretable, biologically relevant sources of expression variation in the compendium.

### iModulon Characterization

To facilitate iModulon characterization, we utilized the PyModulon Python package (Sastry et al., 2021b). As *S. acidocaldarius* has a poorly documented TRN, k-means clustering (with k=3) was utilized to identify component-specific thresholds. This technique clustered genes into 3 groups based on the magnitude of each gene’s weighting. The cluster with the lowest average weighting was filtered out, and its bounds used as the cutoff in determining which genes were part of an iModulon. Each iModulon was then compared to the draft TRN table to find iModulons with significant overlap with known regulons. Next, KEGG and GO annotations were utilized to identify iModulons with significant overlap with known metabolic pathways. The remaining iModulons were functionally mapped by analyzing their activities and literature review. The scripts showing these characterizations can be found at: https://github.com/SBRG/modulome_saci/blob/master/notebooks/5_a_iModulon_characterization_GO_KEGG_setup.ipynb, https://github.com/SBRG/modulome_saci/blob/master/notebooks/5_b_iModulon_characterization_KEGG_enrichments.ipynb, and https://github.com/SBRG/modulome_saci/blob/master/notebooks/5_c_iModulon_characterization_remaining_iModulons.ipynb.

### Generating iModulonDB Dashboards

iModulonDB dashboards were generated using the PyModulon package (Rychel et al., 2021; Sastry et al., 2021b); the pipeline for doing so can be found at https://pymodulon.readthedocs.io/en/latest/tutorials/creating_an_imodulondb_dashboard.html.

## Results

### Independent Component Analysis reveals the structure of the *S. acidocaldarius* transcriptome

To prepare the data compendium, we first compiled all publicly available RNA-seq data available from NCBI Sequence Read Archive (Kodama et al., 2012). After processing the data and filtering through a quality control pipeline (see Methods, **Figure 1A**), the final high-quality compendium consisted of 95 RNA-seq experiments, ranging over a diverse set of conditions, including starvation (time-course), acid stress, and UV irradiation (time-course). Application of ICA to this compendium resulted in 45 robust iModulons, each of which constitutes a statistically independent signal of gene expression. Although these 45 iModulons (**Figure 1D**) only contain 755 genes (32% of the 2351 genes found on the genome), the iModulons can explain 72% of the observed variance in gene expression; genes which were not captured in any iModulons did not vary much in expression. Gene distribution in these iModulons follows a power law, with a few iModulons containing a large number of genes, while most contain fewer than 20 genes (**Supplementary Figure 1**).

In order to characterize the iModulons, we first built a scaffold TRN (**Supplementary File 1**) based on the existing literature for *S. acidocaldarius*, consisting of 346 gene-regulator interactions for 28 regulators (Koerdt et al., 2011; Reimann et al., 2012; Lassak et al., 2013; Märtens et al., 2013; Orell et al., 2013; Ouchi et al., 2013; Vassart et al., 2013; Anjum et al., 2015; Buetti-Dinh et al., 2016; Liu et al., 2016; Haurat et al., 2017; Li et al., 2017; Schult et al., 2018; Wagner et al., 2018; Lemmens et al., 2019; Wang et al., 2019; Baes et al., 2020; Stracke et al., 2020; Suzuki et al., 2020; van der Kolk et al., 2020). To our knowledge, this is the first comprehensive collection of gene-regulator interactions compiled for *S. acidocaldarius*. We used this scaffold to infer which regulon strongly overlapped with each iModulon (See Methods). The iModulon-derived TRN of *S. acidocaldarius* consists of 957 gene-iModulon interactions, of which 65 are known gene-regulator interactions and the remaining 892 are new predictions. These newly predicted interactions provide a roadmap for regulon discovery as well as an opportunity for regulator-agnostic, data-driven discovery of the *S. acidocaldarius* TRN.

### iModulons reveal the basis for the variability in the *S. acidocaldarius* transcriptome

Unlike regulons, which require direct experimental evidence of TF binding sites, iModulons generate sets of co-regulated genes (even those with no known regulator) through extracting global patterns in the transcriptome. The resulting gene sets can be mapped onto known regulons or known cellular or biochemical processes. This curation has allowed for the functional characterization of 37 out of 45 iModulons, despite many gaps in this organism’s TRN. These characterized iModulons, which are clustered into 5 main groupings, account for approximately 48% of the explained variance in the compendium. The smallest grouping (accounting for genomic alterations to specific strains included in the compendium) explains <1% of the variance, while the largest grouping (accounting for other functional aspects of cellular regulation) explains 22% of variance. Interestingly, stress-response iModulons and Vitamin B iModulons both explain a similar percentage of the variance in the compendium (6% and 4% respectively), with miscellaneous metabolic iModulons explaining the remaining 15%.

### Archaeal iModulons Recapitulate Known Regulons in Literature

Of the 45 robust iModulons generated by ICA, 5 recapitulate known regulons in literature: ArnRAB, FadR, YtrA, UV-tfb3, and LysM. An investigation into these iModulons validates that iModulons represent the effects of transcriptional regulators in archaea, and provide additional biological insight into the TRN beyond what currently known regulons can. In the subsequent sections, we provide three specific examples.

### The ArnRAB iModulon recapitulates core archaellum formation genes

The ArnRAB iModulon consists of the core archaellum formation genes of the arl operon (*Saci_1172* to *Saci_1179*, **Figure 2A, 2B**) (Lassak et al., 2013). This operon forms archaeal-specific motility pili that are the primary drivers of cell movement, especially in starvation conditions (**Figure 2C**) (Beeby et al., 2020). The *arl* operon is regulated by a complex interplay of TFs (Reimann et al., 2012; Lassak et al., 2013; Haurat et al., 2017; Ye et al., 2020). The activity of the iModulon also reflect observations from the literature; namely, upregulation over time in response to nutrient-limiting conditions (**Figure 2D**). The activity level of this iModulon is a useful measurement that likely combines the effects of several regulators: high activity could be the result of *arl* operon activation by ArnR/ArnR1, and low activity may be due to repression by ArnA/ArnB.

**Figure 2:**
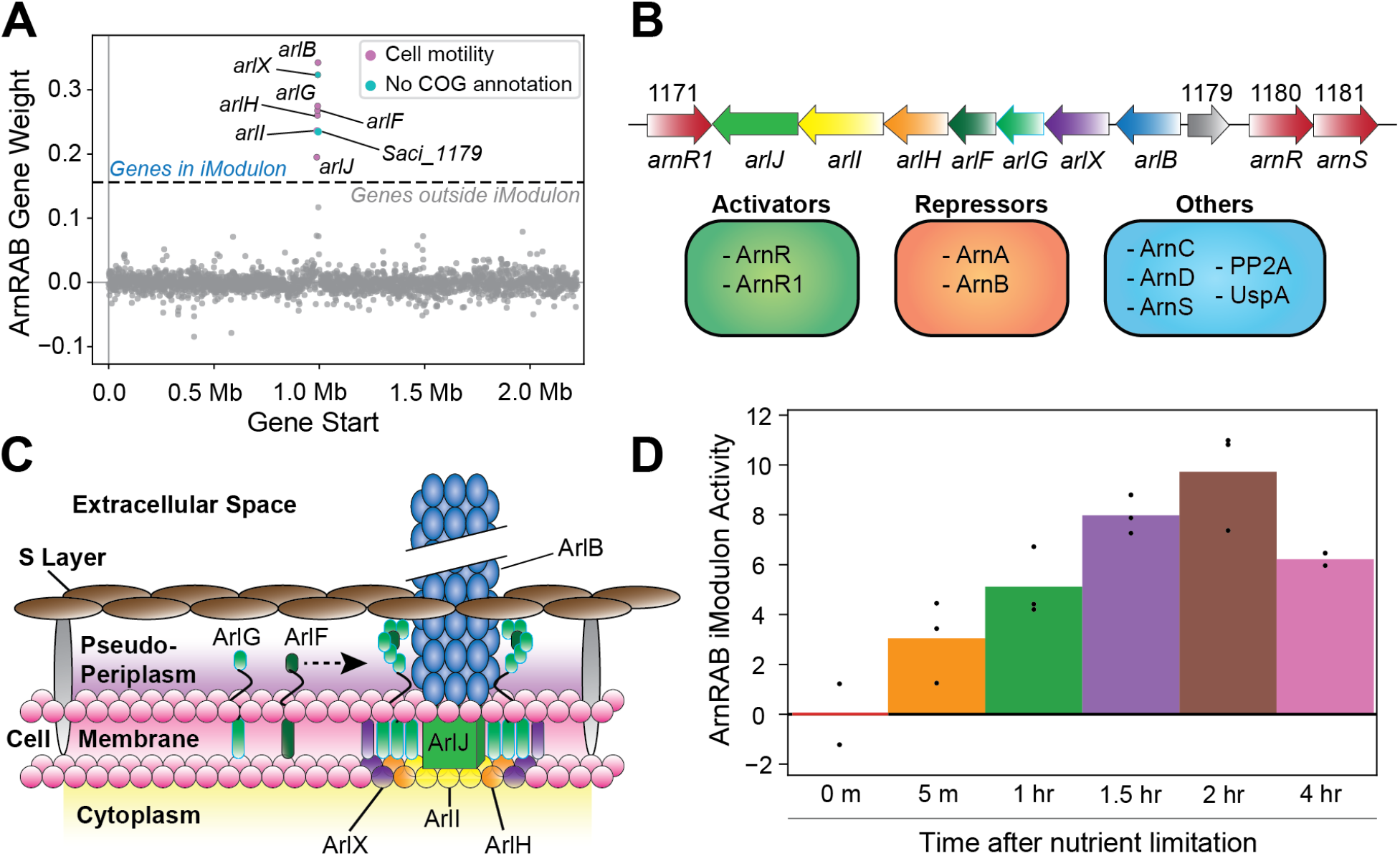
Overview of the ArnRAB iModulon. **A**.) Scatter plot of the gene weights for the ArnRAB iModulon. The x-axis represents the genomic location and the y-axis represents the weight of each gene in the independent component. The dashed lines represent the threshold beyond which genes of the independent component are included in an iModulon. **B**.) Gene map of archaellum formation operon. Known regulatory elements are also listed. **C**.) Structural map showcasing individual gene products from the archaellum formation operon and how each gene product integrates to form these archaeal motility pili. **D**.) Bar plot depicting the ArnRAB iModulon activity over time in nutrient-limiting conditions. Bars represent averages and points represent individual replicate samples. All activity levels are relative to a chosen control condition (0 minutes after nutrient limitation in this case).

### The FadR iModulon successfully captures its corresponding gene cluster

The FadR iModulon represents another case study that demonstrates the ability of ICA to extract co-regulated gene clusters from archaeal RNA-seq compendia. This iModulon (**Supplementary Figure 2**) consists of 19 genes in the FadR_Sa_ gene cluster (*Saci_1103* to *Saci_1122*) (Wang et al., 2019), excluding *fadR* itself (*Saci_1107*), as well as six additional genes not present in the regulon. However, the iModulon is missing five genes found in the regulon: *fadR* and *Saci_1123* to *Saci_1126*. The absence of *fadR* in this iModulon is explained by the fact that *fadR* is contained in its own single-gene iModulon, which captures the FadR knockout condition (FadR-KO). The remaining four genes are regulated by FadR_Sa_ as it binds near *Saci_1123*. However, the confirmed FadR-binding region for this section of the gene cluster is known to be much weaker (Wang et al., 2019), which could result in a weaker signal that could not be detected in this dataset. It is worth noting, however, that *Saci_1126* is just below the statistical threshold for enrichment in this iModulon (0.079 vs 0.08). There are also six new genes in the FadR iModulon that are not in the currently known regulon (**Supplementary Figure 2**). All six genes are functionally uncharacterized, with the exception of *Saci_1992*; this gene is a CRISPR-associated TF (Benninghoff et al., 2021). Five of these genes have negative gene weights, which means that their expression is anti-correlated with the expression of the FadR_Sa_ gene cluster. In this example, the differences between the regulon and iModulon capture differences in binding strength that have been identified previously, as well as proposing new putative FadR-regulated genes.

The FadR iModulon is activated when FadR_Sa_ is knocked-out or during nutrient-limiting conditions (**Supplementary Figure 2**). This suggests derepression in these conditions, as FadR_Sa_ is known to repress its own gene cluster (Wang et al., 2019).

### iModulons suggest an expanded role for LysM

The LysM iModulon recapitulates canonical lysine biosynthesis in *S. acidocaldarius*. The iModulon consists of 10 genes, of which 4 make up the LysM-regulated *lysWXJK* operon, which codes for enzymes in the LysW-mediated α-aminoadipate pathway (Zabriskie and Jackson, 2000; Ouchi et al., 2013; Suzuki et al., 2020). **(Figure 3A**). Incidentally, the *lysYZM* operon, which also codes for two enzymes (LysY and LysZ) in the α-aminoadipate pathway, is absent from this iModulon. This absence suggests that some other regulatory elements may also influence the expression of *lysYZM*, and thus the entire LysM iModulon indirectly (**Figure 3B**).

**Figure 3:**
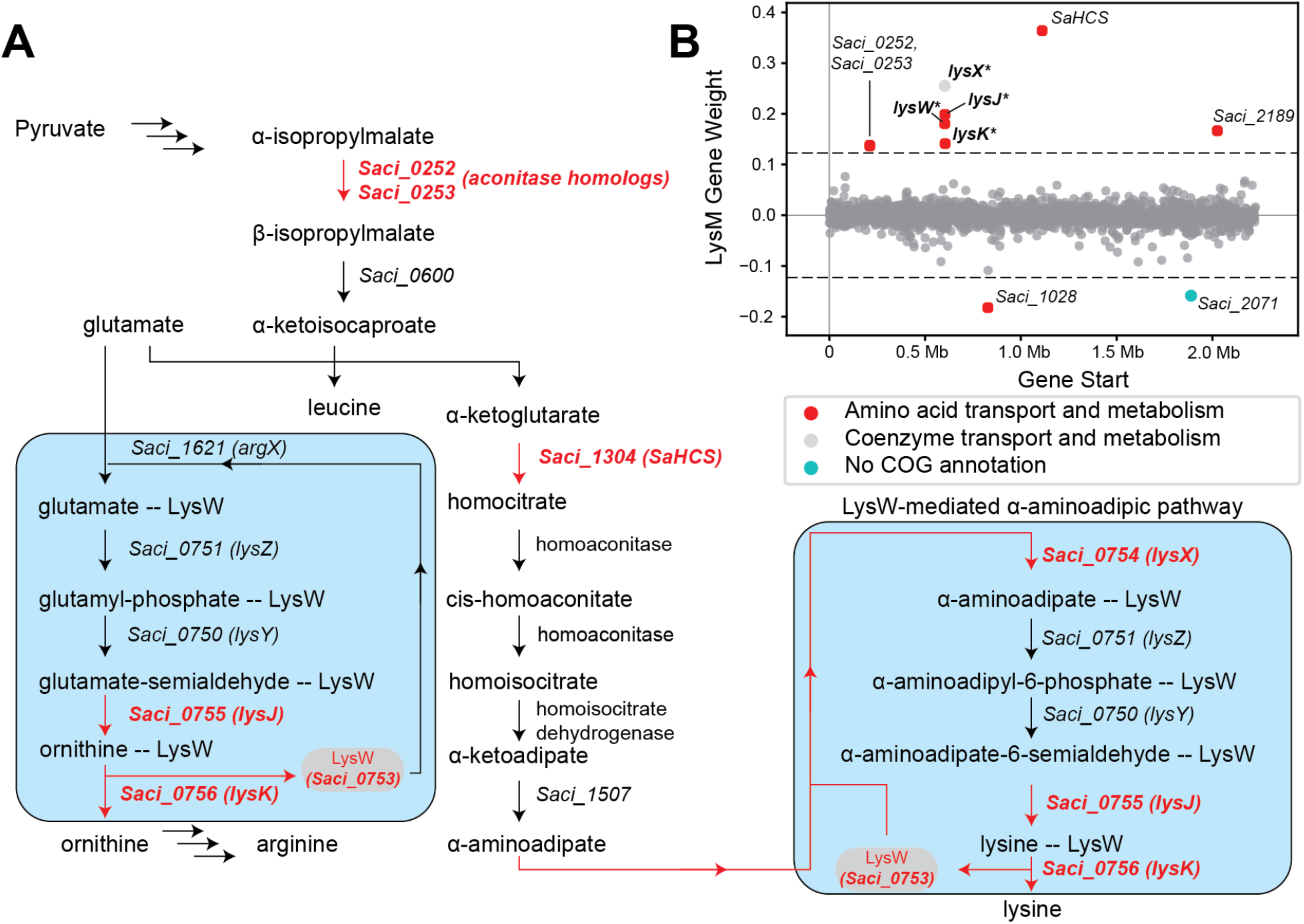
Overview of the LysM regulon and iModulon. **A**.) Pathway map overviewing lysine biosynthesis in *S. acidocaldarius*. Genes in red are part of the LysM iModulon. The blue boxes specifically show the LysW-mediated pathway used in lysine and arginine biosynthesis which consists of *lysYZMWXKJ*. LysW attaches to the amino group of glutamate/ɑ-aminoadipate until the substrate forms into ornithine/lysine, respectively **B**.) Scatter plot of the gene weights for the LysM iModulon. Genes are colored by COG categories.

Additionally, this iModulon contains *Saci_1304*, which codes for homocitrate synthase (SaHCS) (Suzuki et al., 2020); this enzyme catalyzes the first step towards lysine biosynthesis in *S. acidocaldarius* (Ouchi et al., 2013; Suzuki et al., 2020). *The LysM iModulon also contains Saci_0252* and *Saci_0253*, which encode for 3-isopropylmalate dehydratase, a part of the leucine biosynthesis pathway (Park et al., 11/2018; Quehenberger et al., 2019). These genes are also aconitase homologs (Gross et al., 1963; Calvo et al., 1964; Cole et al., 1973), and may also have catalytic activity in homocitrate to homoisocitrate conversion (**Figure 3B**).

Finally, this iModulon consists of 3 poorly characterized genes: *Saci_1028, Saci_2071*, and *Saci_2189. Saci_1028* putatively codes for an acetyl-ornithine aminotransferase family protein and *Saci_2189* putatively codes for an APC family permease (potentially an aspartate-proton symporter), while *Saci_2071* is completely uncharacterized, encoding a hypothetical protein. It should be noted that *Saci_1028* and *Saci_2071* are the only genes in this iModulon to have negative gene weights. This indicates that the expression of both of these genes are anti-correlated with the rest of the genes in this iModulon; that is, these genes are upregulated when the rest of the iModulon is downregulated and vice versa. The presence of these genes in the LysM iModulon indicates that they are likely involved in lysine biosynthesis, and perhaps regulated by LysM or some other shared regulatory mechanism.

However, LysM, a TF which is highly conserved in *Sulfolobus* species, is known to bind with at least 10 different amino acid effectors in the closely-related species *Saccharolobus solfataricus* (previously *Sulfolobus solfataricus*) (Song et al., 2013-3). This affinity with multiple amino-acid effectors could reasonably extend to the homologous LysM TF in *S. acidocaldarius*. This fact, combined with the proven ability of ICA to extract regulatory signals, leads us to hypothesize that all 10 genes of this iModulon are regulated by LysM. Alternatively, both the *lysYZM* and *lysWXKJ* operons, alongside the remaining enriched genes of this iModulon, could be regulated together by a common regulatory element(s). In either case, iModulons direct us towards 6 genes which warrant additional investigation, and may further elucidate amino acid biochemistry and its associated TRN in *S. acidocaldarius*.

### iModulons uncover data-driven targets for gene function and regulator discovery

As seen with LysM, iModulons also function as data-driven aids for gene/regulator discovery. Rather than searching for regulators and their known binding locations along a genome, iModulons provide a set of co-regulated genes, and instead allow for a data-driven discovery of common regulatory elements. The following sections provide three such examples for the *S. acidocaldarius* TRN.

### Uncharacterized genes may compose the UV-induced DNA export system

The UV-tfb3 iModulon contains genes that form the *tfb3*-dependent UV stress response (**Figure 4A**) (Schult et al., 2018). Under UV irradiation, *S. acidocaldarius* cells aggregate, form specific adhesion pili, and exchange DNA with each other to repair their genomes (Götz et al., 2007; Schult et al., 2018). This process is performed in a *tfb3*-dependent manner, which is also reflected by this iModulon’s activities: steadily increasing after UV irradiation in wild-type cells, but almost completely absent in *tfb3*-disrupted mutants (**Figure 4B**). Two systems have been identified that help compose this response: the *ups* operon and the *ced* system. The *ups* operon (*upsXEFAB*) (Ajon et al., 2011; van Wolferen et al., 2013, 2015) consists of genes that code for the specific pili that aid *S. acidocaldarius* cells in adhering to each other post-UV irradiation. The *ced* system (*cedA/A1/A2/B*) (van Wolferen et al., 2016) consists of multiple transporters that import DNA. However, a crucial aspect of this response is still unidentified: the DNA export system. As the *ced* system only imports DNA, a corresponding DNA export system must exist for *S. acidocaldarius* cells to properly exchange DNA.

**Figure 4:**
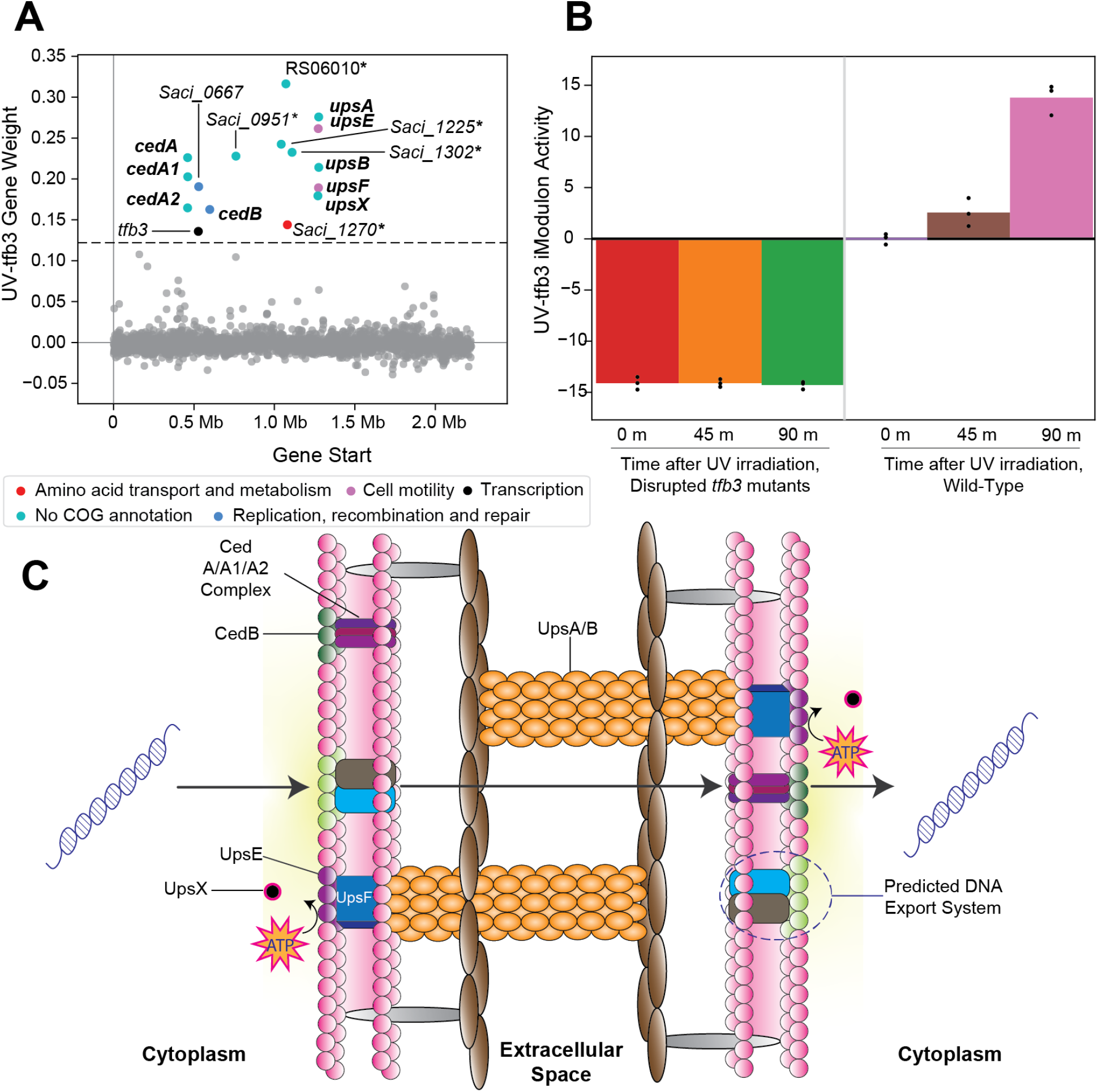
Overview of the *tfb3*-dependent UV-stress response iModulon (UV-tfb3). **a**.) Scatter plot of the UV-tfb3 iModulon gene weights. Genes are colored by COG categories. Genes in bold are part of the known *tfb3*-dependent response. Genes with an asterisk are proposed to make up the DNA export system. **b**.) UV-tfb3 iModulon activity plotted over time after UV-irradiation. The left three bars show the activities of the *tfb3* insertion mutants (relative to wild-type cells before UV irradiation). The right three bars show the activities of the wild-type cells (relative to wild-type cells before UV irradiation). **c**.) A visual overlay of the *tfb3*-dependent UV-stress response.

The UV-tfb3 iModulon, in addition to the *ups* operon and the *ced* system, contains five uncharacterized genes, which we propose to constitute this undiscovered DNA export system (**Figure 4C**). InterPro scans of all the uncharacterized genes revealed that *Saci_1270* and

*Saci_1302* contained multiple transmembrane, cytosolic, and non-cytosolic domains, providing further evidence that these genes encode enzymes that help export DNA (**Supplementary File 3**).

### iModulons provide evidence of an uncharacterized global regulator

ICA decomposition of our RNA-seq compendium revealed 45 iModulons, which we ranked by explained variance in Figure 1D. iModulons with high explained variance are nearly always regulated by global transcriptional regulators (Lamoureux et al., 2021). However, the iModulon which explained the largest variance in the *S. acidocaldarius* data was primarily enriched with uncharacterized or poorly characterized genes. This iModulon consists of 33 enriched genes: 11 characterized genes, 12 poorly characterized genes, and 10 completely uncharacterized genes (**Figure 5A**). Of the genes which are characterized, six genes encode acyl-CoA ligases, and of the genes which are poorly characterized, five are membrane-bound proteins. Beyond this information, however, little is known about the enriched genes of this iModulon.

**Figure 5:**
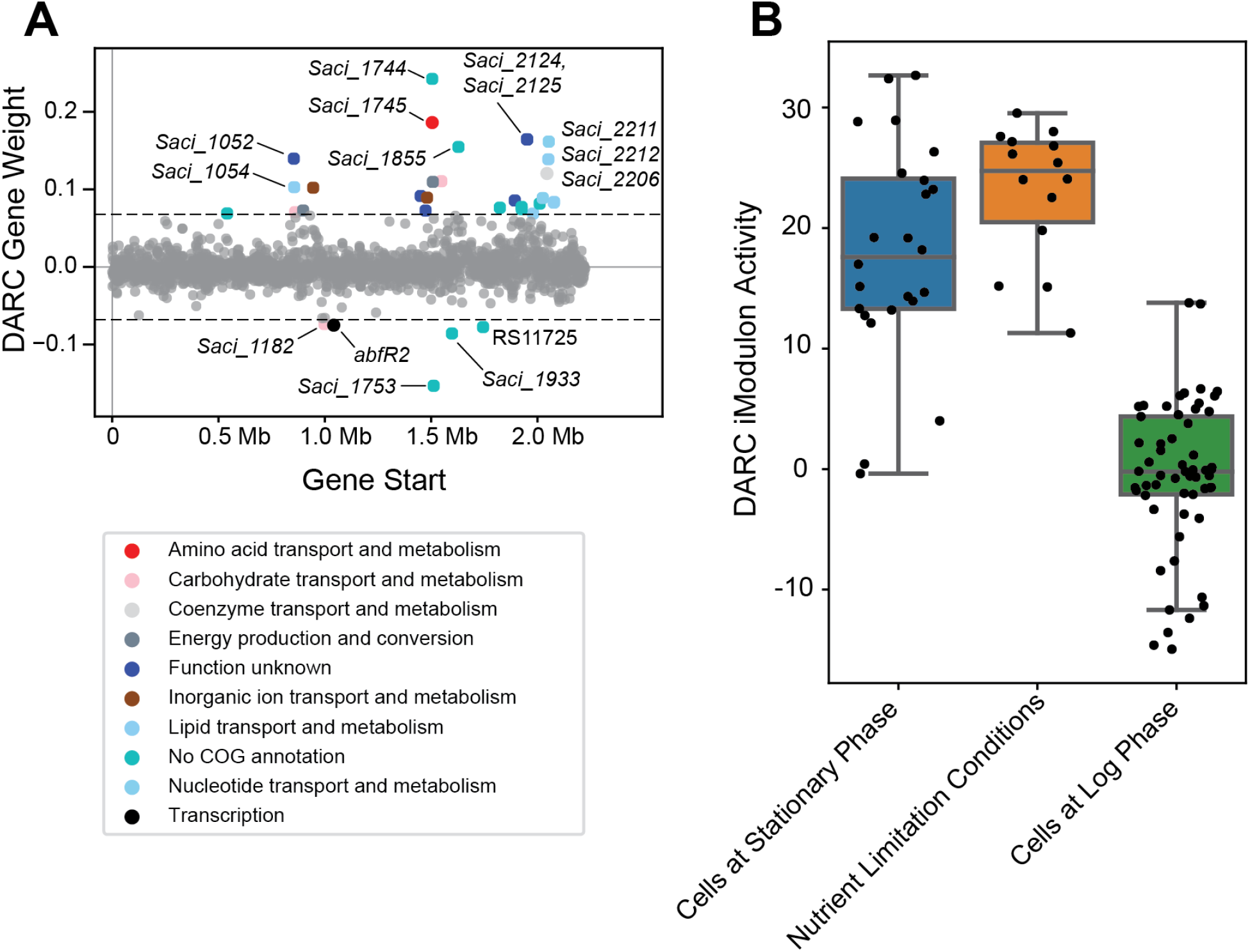
Overview of the “DARC” iModulon. **a**.) Scatter plot of the gene weightings for this iModulon. Genes are colored by COG categories. **b**.) Boxplot showcasing this iModulon’s activities for various conditions. Dots represent individual sample values. “Cells at stationary phase” includes all samples in the RNA-seq compendium which were collected for cells in stationary phase. “Cells at log phase” includes all samples in the RNA-seq compendium which were collected for cells in log phase, except for cells which were exposed to any nutrient-limitation conditions; “Nutrient Limitation Conditions” represents those such samples.

The largest difference in this iModulon’s activity is between cells grown in log phase and cells grown in stationary phase or in nutrient-limiting conditions (**Figure 5B**). This iModulon contains two TFs: *Saci_1223* (*abfR2*), a secondary biofilm regulator, and *Saci_2103*, a predicted MarR family TF. Taken together, we propose that this iModulon is related to the cell membrane, possibly in a growth-dependent manner, but further work is needed to fully elucidate the role of this iModulon in the overall TRN of *S. acidocaldarius*. As this iModulon represents a biological signal (extracted by ICA), but with no known regulators or clearly defined function, we name this iModulon a Discovered signal with Absent Regulatory Components, or DARC.

### The XylR-SoxM iModulon may be governed by a global regulator and could contain an undiscovered, peptide-induced sugar transporter

The XylR-SoxM iModulon consists of 47 enriched genes (**Figure 6A**), of which 10 are part of the XylR regulon (Wagner et al., 2018; van der Kolk et al., 2020), and 6 of the *soxEFGHIM* gene cluster (Park et al., 11/2018). This iModulon also contains *Saci_2032, Saci_2033*, and *Saci_2034*, which are all genes predicted to code for enzymes in glycerol uptake and metabolism. The remaining enriched genes are poorly characterized, and mostly encode either thiolases, thioredoxins, or hypothetical proteins. Of the 10 genes shared between the XylR regulon and the XylR-SoxM iModulon (**Figure 6B**), 8 genes are known to be downregulated in xylose-growth conditions: *Saci_1147, Saci_1148*, and *Saci_2230* to *Saci_2235* (Wagner et al., *2018). The remaining 2, which are upregulated in xylose-growth conditions, are Saci_2122* (*xylF*) and *Saci_1760. Saci_1760* codes for a glycosylated membrane protein which is only present during D-xylose/L-arabinose growth conditions, while *Saci_2122* encodes a D-xylose/L-arabinose substrate-binding protein which works in concert with transport proteins XylG and XylH to import pentose sugars (Wagner et al., 2018; van der Kolk et al., 2020). However, *xylG* and *xylH* are not enriched in this iModulon.

**Figure 6:**
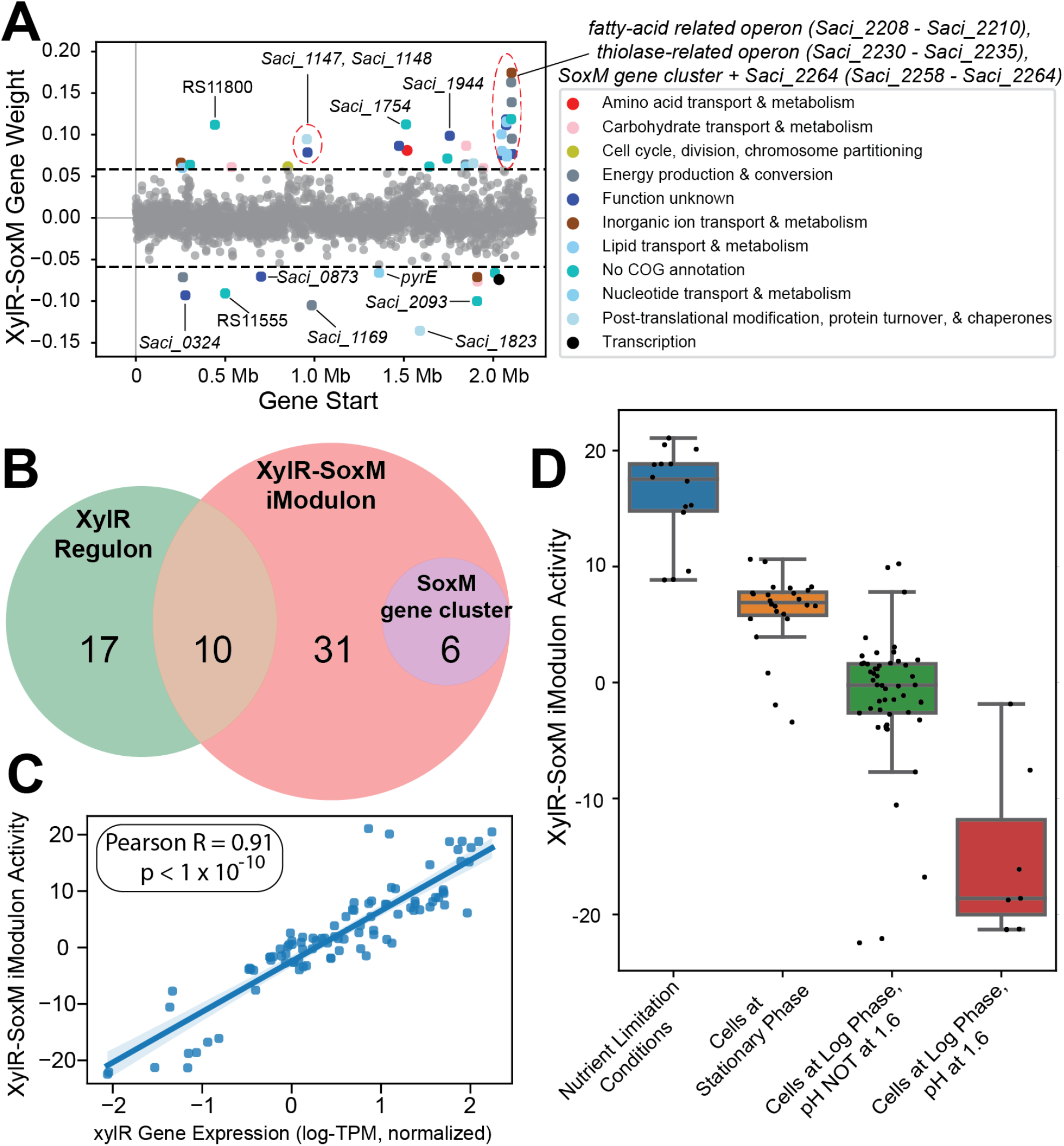
Overview of the XylR-SoxM iModulon. **A**.) Scatter plot of the gene weightings for this iModulon. Genes are colored by COG categories. **B**.) Venn diagram comparing the XylR regulon, XylR-SoxM iModulon, and the SoxM gene cluster. **C**.) Scatterplot of *xylR* gene expression (x-axis) vs. XylR-SoxM iModulon activity (y-axis). The correlation coefficient for this data is 0.91. **D**.) Boxplot of XylR-SoxM activities by condition. “Nutrient Limitation Conditions” refers to all samples in the RNA-seq compendium which were exposed to any nutrient-limitation conditions.

Taken together, this information suggests that pentose sugar uptake may have at least two forms of regulation. The first, which is captured by this iModulon, consists of the regulation of various genes, including *Saci_2122, Saci_1760*, and the entire SoxM gene cluster (which consists of an archaeal cytochrome system) (Komorowski et al., 2002). The second, which is not captured by this iModulon, should consist of the regulation of the remainder of the XylR regulon (*xylG, xylH*, etc.) and possibly also the genes involved in the aldolase-independent Weimberg pathway, responsible for converting D-xylose/L-arabinose into ɑ-ketoglutarate (Wagner et al., 2018; van der Kolk et al., 2020). Having multiple regulatory substructures agrees with previous assertions which state that there may be additional layers of regulation in pentose sugar uptake and metabolism, beyond just the regulator XylR (Wagner et al., 2018; van der Kolk et al., 2020). Interestingly, while *Saci_2116* (*xylR*) is not enriched in this iModulon, it is just below the statistical threshold (0.057 vs 0.058) and its expression is highly correlated with this iModulon’s activity (Pearson-R = 0.91, **Figure 6C**). This iModulon also has positive activity in nutrient-limiting conditions and in stationary phase, with negative activity in log phase (**Figure 6D**), further suggesting that this iModulon contains genes related to growth and/or starvation. Altogether, this suggests two potential hypotheses: (1) *xylR* is a global regulator that has an expanded function related to growth, or (2) there is a separate global regulator that regulates all the genes in this iModulon (and potentially also *xylR*). To test these hypotheses, more RNA-seq data must be accumulated for *S. acidocaldarius*.

Additionally, this iModulon may include a gene which encodes an as-yet-undiscovered xylose transporter in *S. acidocaldarius*. While *xylG* and *xylH* both form known D-xylose transporters, there is evidence for the existence of another mechanism which can transport D-xylose into *S. acidocaldarius* in the presence of peptides (Wagner et al., 2018), but the gene which may encode such a transporter is currently unknown. There are three genes in the XylR-SoxM iModulon that encode putative transporters: *Saci_0324, Saci_0675*, and *Saci_2095*. In particular, *Saci_2095* is predicted to encode an MFS transporter (likely a sugar transporter). Additionally, the EggNog cluster of orthologous genes (COG) mapper annotates this gene’s function as carbohydrate transport and metabolism. Furthermore, an InterPro scan was performed on this gene’s amino acid sequence and resulted in this gene being classified as a sugar transporter (**Supplementary File 4**).

In summary, our investigation of this iModulon resulted in evidence suggesting regulation by a global regulator, as well as the potential identification of *Saci_2095* as a peptide-induced D-xylose transporter.

## Discussion

Here, we collated all publicly available data to generate a high-quality RNA-seq compendium on *S. acidocaldarius*, and deconvoluted it using ICA. This deconvolution extracted 45 iModulons, whose overall activity can explain 72% of the variance in gene expression across the wide range of conditions contained in the compendium. 37 of these iModulons correspond to specific biological functions, with 5 corresponding to known regulators. We analyzed the enriched gene sets of these iModulons and presented findings that agree with previously existing knowledge. In addition, we generated data-driven hypotheses that could be experimentally tested in future investigations. We also showcase a previously unknown set of co-regulated genes which form an uncharacterized iModulon that explains 12% of the variance in the compendium. These results also generate the first comprehensive global TRN structure of *S. acidocaldarius*.

Through ICA, we identified well-studied regulons with high accuracy (such as ArnRAB, FadR, UV-tfb3 iModulons), in addition to discovering much more of the TRN landscape in *S. acidocaldarius*. ICA detected potential new regulatory targets for LysM, which may also further elucidate amino acid metabolism in *S. acidocaldarius*. Similarly, the added presence of 5 uncharacterized genes in the UV-tfb3 iModulon suggests more regulatory targets for *tfb3*, as they may form the currently undiscovered DNA export system for this organism. Completely new sections of TRN are also unearthed by ICA, as shown by the observation that the 33 co-regulated genes in the DARC iModulon account for 12% of the total variance in the compendium, by far the largest of any iModulon. Many of these genes are poorly understood, and warrant further investigation. The ability of iModulons to identify novel regulatory signals enables improved data-driven discovery over traditional differential gene expression studies.

Beyond the static plots provided in this manuscript, all generated iModulon data and interactive graphical summaries are available for interrogation online at iModulonDB.org (Rychel et al., 2021). Code for the analysis pipeline used is hosted on GitHub (https://github.com/SBRG/modulome_saci/).

As with many other machine learning methods, the results generated from ICA will improve as more high-quality data is used to generate the initial RNA-seq compendium. Already, many recently published RNA-seq studies for *S. acidocaldarius* exist which were not included in the compendium, as the data was unavailable at the onset of this project. Addition of such high-quality data, combined with transcriptome analysis under further unique conditions, will allow for a higher-resolution insight into the TRN. Larger, multifunction iModulons will likely separate into more biologically accurate modules. This division may also capture new regulons, some which may recapitulate existing knowledge of known regulators (e.g., BarR, Sa-Lrp, AbfR1) and others which may reveal further insights. In principle, if transcriptomic data for every possible condition were to be obtained for this organism, ICA would generate a comprehensive, data-driven, and quantitatively-irreducible TRN.

## Supporting information

Supplementary Information (Main)

Supplementary File 1 (draft TRN)

Supplementary File 2 (gene annot)

Supplementary File 3 (InterPro UV)

Supplementary File 4 (InterPro XylR)

## Acknowledgements

The authors would like to acknowledge Dr. Amitesh Anand, Dr. Sonja-Verena Albers, and Dr. Marleen van Wolferen for useful discussions.

## References

Ajon, M., Fröls, S., van Wolferen, M., Stoecker, K., Teichmann, D., Driessen, A. J. M., et al. (2011) UV-inducible DNA exchange in hyperthermophilic archaea mediated by type IV pili. Mol. Microbiol. 82, 807–817. doi:10.1111/j.1365-2958.2011.07861.x.

Albers, S.-V., and Meyer, B. H. (2011) The archaeal cell envelope. Nat. Rev. Microbiol. 9, 414–426. doi:10.1038/nrmicro2576.

Anjum, R. S., Bray, S. M., Blackwood, J. K., Kilkenny, M. L., Coelho, M. A., Foster, B. M., et al. (2015) Involvement of a eukaryotic-like ubiquitin-related modifier in the proteasome pathway of the archaeon Sulfolobus acidocaldarius. Nat. Commun. 6, 8163. doi:10.1038/ncomms9163.

Baes, R., Lemmens, L., Mignon, K., Carlier, M., and Peeters, E. (2020) Defining heat shock response for the thermoacidophilic model crenarchaeon Sulfolobus acidocaldarius. Extremophiles 24, 681–692. doi:10.1007/s00792-020-01184-y.

Beeby, M., Ferreira, J. L., Tripp, P., Albers, S.-V., and Mitchell, D. R. (2020) Propulsive nanomachines: the convergent evolution of archaella, flagella and cilia. FEMS Microbiol. Rev 44, 253–304. doi:10.1093/femsre/fuaa006.

Benninghoff, J. C., Kuschmierz, L., Zhou, X., Albersmeier, A., Pham, T. K., Busche, T., et al. (2021) Response of the thermoacidophilic Archaeon Sulfolobus acidocaldarius to solvent stress exemplified by 1-butanol exposure. Appl. Environ. Microbiol. doi:10.1128/AEM.02988-20.

Biton, A., Bernard-Pierrot, I., Lou, Y., Krucker, C., Chapeaublanc, E., Rubio-Pérez, C., et al. (2014) Independent component analysis uncovers the landscape of the bladder tumor transcriptome and reveals insights into luminal and basal subtypes. Cell Rep. 9, 1235–1245. doi:10.1016/j.celrep.2014.10.035.

Brock, T. D., Brock, K. M., Belly, R. T., and Weiss, R. L. (1972) Sulfolobus: a new genus of sulfur-oxidizing bacteria living at low pH and high temperature. Arch. Mikrobiol. 84, 54–68. doi:10.1007/BF00408082.

Buetti-Dinh, A., Dethlefsen, O., Friedman, R., and Dopson, M. (2016) Transcriptomic analysis reveals how a lack of potassium ions increases Sulfolobus acidocaldarius sensitivity to pH changes. Microbiology 162, 1422–1434. doi:10.1099/mic.0.000314.

Calvo, J. M., Stevens, C. M., Kalyanpur, M. G., and Umbarger, H. E. (1964) THE ABSOLUTE CONFIGURATION OF ALPHA-HYDROXY-BETA-CARBOXYISOCAPROIC ACID (3-ISOPROPYLMALIC ACID), AN INTERMEDIATE IN LEUCINE BIOSYNTHESIS. Biochemistry 3, 2024–2027. doi:10.1021/bi00900a043.

Cole, F. E., Kalyanpur, M. G., and Stevens, C. M. (1973) Absolute configuration of alpha isopropylmalate and the mechanism of its conversion to beta isopropylmalate in the biosynthesis of leucine. Biochemistry 12, 3346–3350. doi:10.1021/bi00741a031.

Di Tommaso, P., Chatzou, M., Floden, E. W., Barja, P. P., Palumbo, E., and Notredame, C. (2017) Nextflow enables reproducible computational workflows. Nat. Biotechnol. 35, 316–319. doi:10.1038/nbt.3820.

Ewels, P., Magnusson, M., Lundin, S., and Käller, M. (2016) MultiQC: summarize analysis results for multiple tools and samples in a single report. Bioinformatics 32, 3047–3048. doi:10.1093/bioinformatics/btw354.

Götz, D., Paytubi, S., Munro, S., Lundgren, M., Bernander, R., and White, M. F. (2007) Responses of hyperthermophilic crenarchaea to UV irradiation. Genome Biol. 8, R220. doi:10.1186/gb-2007-8-10-r220.

Gross, S. R., Burns, R. O., and Umbarger, H. E. (1963) THE BIOSYNTHESIS OF LEUCINE. II. THE ENZYMIC ISOMERIZATION OF BETA-CARBOXY-BETA-HYDROXYISOCAPROATE AND ALPHA-HYDROXY-BETA-CARBOXYISOCAPROATE. Biochemistry 2, 1046–1052. doi:10.1021/bi00905a023.

Haurat, M. F., Figueiredo, A. S., Hoffmann, L., Li, L., Herr, K.J, Wilson, A., et al. (2017) ArnS, a kinase involved in starvation-induced archaellum expression. Mol. Microbiol. 103, 181–194. doi:10.1111/mmi.13550.

Hyvärinen, A., and Oja, E. (2000) Independent component analysis: algorithms and applications. Neural Netw. 13, 411–430. doi:10.1016/s0893-6080(00)00026-5.

IOS Press Ebooks - Jupyter Notebooks – a publishing format for reproducible computational workflows Available at: https://ebooks.iospress.nl/publication/42900 [Accessed March 12, 2021].

Jutten, C., and Herault, J. (1991) Blind separation of sources, part I: An adaptive algorithm based on neuromimetic architecture. Signal Processing 24, 1–10. doi:10.1016/0165-1684(91)90079-X.

Kodama, Y., Shumway, M., Leinonen, R., and International Nucleotide Sequence Database Collaboration (2012). The Sequence Read Archive: explosive growth of sequencing data. Nucleic Acids Res. 40, D54–6. doi:10.1093/nar/gkr854.

Koerdt, A., Orell, A., Pham, T. K., Mukherjee, J., Wlodkowski, A., Karunakaran, E., et al. (2011) Macromolecular fingerprinting of sulfolobus species in biofilm: a transcriptomic and proteomic approach combined with spectroscopic analysis. J. Proteome Res. 10, 4105–4119. doi:10.1021/pr2003006.

Komorowski, L., Verheyen, W., and Schäfer, G. (2002) The archaeal respiratory supercomplex SoxM from S. acidocaldarius combines features of quinole and cytochrome c oxidases. Biol. Chem. 383, 1791–1799. doi:10.1515/BC.2002.200.

Lamoureux, C. R., Decker, K. T., Sastry, A. V., McConn, J. L., Gao, Y., and Palsson, B. O. (2021) PRECISE 2.0 - an expanded high-quality RNA-seq compendium for Escherichia coli K-12 reveals high-resolution transcriptional regulatory structure. bioRxiv, 2021.04.08.439047. doi:10.1101/2021.04.08.439047.

Langmead, B., Trapnell, C., Pop, M., and Salzberg, S. L. (2009) Ultrafast and memory-efficient alignment of short DNA sequences to the human genome. Genome Biol. 10, R25. doi:10.1186/gb-2009-10-3-r25.

Lassak, K., Peeters, E., Wróbel, S., and Albers, S.-V. (2013) The one-component system ArnR: a membrane-bound activator of the crenarchaeal archaellum. Mol. Microbiol. 88, 125–139. doi:10.1111/mmi.12173.

Lemmens, L., Tilleman, L., De Koning, E., Valegård, K., Lindås, A.-C., Van Nieuwerburgh, F., et al. (2019) YtrASa, a GntR-Family Transcription Factor, Represses Two Genetic Loci Encoding Membrane Proteins in Sulfolobus acidocaldarius. Front. Microbiol. 10, 2084. doi:10.3389/fmicb.2019.02084.

Lewis, A. M., Recalde, A., Bräsen, C., Counts, J. A., Nussbaum, P., Bost, J., et al. (2021) The biology of thermoacidophilic archaea from the order Sulfolobales. FEMS Microbiol. Rev. doi:10.1093/femsre/fuaa063.

Liao, Y., Smyth, G. K., and Shi, W. (2014) featureCounts: an efficient general purpose program for assigning sequence reads to genomic features. Bioinformatics 30, 923–930. doi:10.1093/bioinformatics/btt656.

Li, L., Banerjee, A., Bischof, L. F., Maklad, H. R., Hoffmann, L., Henche, A.-L., et al. (2017) Wing phosphorylation is a major functional determinant of the Lrs14-type biofilm and motility regulator AbfR1 in Sulfolobus acidocaldarius. Mol. Microbiol. 105, 777–793. doi:10.1111/mmi.13735.

Littlechild, J. A. (2015) Archaeal Enzymes and Applications in Industrial Biocatalysts. Archaea 2015, 147671. doi:10.1155/2015/147671.

Liu, H., Wang, K., Lindås, A.-C., and Peeters, E. (2016) The genome-scale DNA-binding profile of BarR, a β-alanine responsive transcription factor in the archaeon Sulfolobus acidocaldarius. BMC Genomics 17, 569. doi:10.1186/s12864-016-2890-0.

Märtens, B., Amman, F., Manoharadas, S., Zeichen, L., Orell, A., Albers, S.-V., et al. (2013) Alterations of the Transcriptome of Sulfolobus acidocaldarius by Exoribonuclease aCPSF2. PLoS One 8, e76569. doi:10.1371/journal.pone.0076569.

McConn, J. L., Lamoureux, C. R., Poudel, S., Palsson, B. O., and Sastry, A. V. (2021) Optimal dimensionality selection for independent component analysis of transcriptomic data. bioRxiv, 2021.05.26.445885. doi:10.1101/2021.05.26.445885.

Orell, A., Peeters, E., Vassen, V., Jachlewski, S., Schalles, S., Siebers, B., et al. (2013) Lrs14 transcriptional regulators influence biofilm formation and cell motility of Crenarchaea. ISME J 7, 1886–1898. doi:10.1038/ismej.2013.68.

Ouchi, T., Tomita, T., Horie, A., Yoshida, A., Takahashi, K., Nishida, H., et al. (2013) Lysine and arginine biosyntheses mediated by a common carrier protein in Sulfolobus. Nat. Chem. Biol. 9, 277–283. doi:10.1038/nchembio.1200.

Park, J., Lee, A., Lee, H.-H., Park, I., Seo, Y.-S., and Cha, J. (11/2018). Profiling of glucose-induced transcription in Sulfolobus acidocaldarius DSM 639. Genes Genomics 40, 1157–1167. doi:10.1007/s13258-018-0675-3.

Poudel, S., Tsunemoto, H., Seif, Y., Sastry, A. V., Szubin, R., Xu, S., et al. (2020) Revealing 29 sets of independently modulated genes in Staphylococcus aureus, their regulators, and role in key physiological response. Proc. Natl. Acad. Sci. U. S. A. 117, 17228–17239. doi:10.1073/pnas.2008413117.

Quehenberger, J., Albersmeier, A., Glatzel, H., Hackl, M., Kalinowski, J., and Spadiut, O. (2019) A defined cultivation medium for Sulfolobus acidocaldarius. J. Biotechnol. 301, 56–67. doi:10.1016/j.jbiotec.2019.04.028.

Quehenberger, J., Shen, L., Albers, S.-V., Siebers, B., and Spadiut, O. (2017) Sulfolobus - A Potential Key Organism in Future Biotechnology. Front. Microbiol. 8, 2474. doi:10.3389/fmicb.2017.02474.

Reimann, J., Lassak, K., Khadouma, S., Ettema, T. J. G., Yang, N., Driessen, A. J. M., et al. (2012) Regulation of archaella expression by the FHA and von Willebrand domain-containing proteins ArnA and ArnB in Sulfolobus acidocaldarius. Mol. Microbiol. 86, 24–36. doi:10.1111/j.1365-2958.2012.08186.x.

Rodionova, I. A., Gao, Y., Sastry, A., Monk, J., Wong, N., Szubin, R., et al. (2020) PtrR (YneJ) is a novel E. coli transcription factor regulating the putrescine stress response and glutamate utilization. bioRxiv, 2020.04.27.065417. doi:10.1101/2020.04.27.065417.

Rodionova, I., Gao, Y., Sastry, A., Hefner, Y., Yoo, R., Rodionov, D., et al. (2021) A novel transcription factor PunR and Nac are involved in purine and purine nucleoside transporter punC regulation in E. coli. Research Square. doi:10.21203/rs.3.rs-146218/v1.

Rychel, K., Decker, K., Sastry, A. V., Phaneuf, P. V., Poudel, S., and Palsson, B. O. (2021) iModulonDB: a knowledgebase of microbial transcriptional regulation derived from machine learning. Nucleic Acids Res. 49, D112–D120. doi:10.1093/nar/gkaa810.

Rychel, K., Sastry, A. V., and Palsson, B. O. (2020) Machine learning uncovers independently regulated modules in the Bacillus subtilis transcriptome. Nat. Commun. 11, 6338. doi:10.1038/s41467-020-20153-9.

Sastry, A. V., Gao, Y., Szubin, R., Hefner, Y., Xu, S., Kim, D., et al. (2019) The Escherichia coli transcriptome mostly consists of independently regulated modules. Nat. Commun. 10, 5536. doi:10.1038/s41467-019-13483-w.

Sastry, A. V., Hu, A., Heckmann, D., Poudel, S., Kavvas, E., and Palsson, B. O. (2021a). Independent component analysis recovers consistent regulatory signals from disparate datasets. PLoS Comput. Biol. 17, e1008647. doi:10.1371/journal.pcbi.1008647.

Sastry, A. V., Poudel, S., Rychel, K., Yoo, R., Lamoureux, C. R., Chauhan, S., et al. (2021b). Mining all publicly available expression data to compute dynamic microbial transcriptional regulatory networks. bioRxiv, 2021.07.01.450581. doi:10.1101/2021.07.01.450581.

Schult, F., Le, T. N., Albersmeier, A., Rauch, B., Blumenkamp, P., van der Does, C., et al. (2018) Effect of UV irradiation on Sulfolobus acidocaldarius and involvement of the general transcription factor TFB3 in the early UV response. Nucleic Acids Res. 46, 7179–7192. doi:10.1093/nar/gky527.

Seemann, T. (2014) Prokka: rapid prokaryotic genome annotation. Bioinformatics 30, 2068–2069. doi:10.1093/bioinformatics/btu153.

Sompairac, N., Nazarov, P. V., Czerwinska, U., Cantini, L., Biton, A., Molkenov, A., et al. (2019) Independent Component Analysis for Unraveling the Complexity of Cancer Omics Datasets. Int. J. Mol. Sci. 20. doi:10.3390/ijms20184414.

Song, N., Nguyen Duc, T., van Oeffelen, L., Muyldermans, S., Peeters, E., and Charlier, D. (2013-3). Expanded target and cofactor repertoire for the transcriptional activator LysM from Sulfolobus. Nucleic Acids Res. 41, 2932–2949. doi:10.1093/nar/gkt021.

Stracke, C., Meyer, B. H., Hagemann, A., Jo, E., Lee, A., Albers, S.-V., et al. (2020) Salt Stress Response of Sulfolobus acidocaldarius Involves Complex Trehalose Metabolism Utilizing a Novel Trehalose-6-Phosphate Synthase (TPS)/Trehalose-6-Phosphate Phosphatase (TPP) Pathway. Appl. Environ. Microbiol. 86. doi:10.1128/AEM.01565-20.

Suzuki, T., Akiyama, N., Yoshida, A., Tomita, T., Lassak, K., Haurat, M. F., et al. (2020) Biochemical characterization of archaeal homocitrate synthase from Sulfolobus acidocaldarius. FEBS Lett. 594, 126–134. doi:10.1002/1873-3468.13550.

Teschendorff, A. E., Journée, M., Absil, P. A., Sepulchre, R., and Caldas, C. (2007) Elucidating the altered transcriptional programs in breast cancer using independent component analysis. PLoS Comput. Biol. 3, e161. doi:10.1371/journal.pcbi.0030161.

van der Kolk, N., Wagner, A., Wagner, M., Waßmer, B., Siebers, B., and Albers, S.-V. (2020) Identification of XylR, the Activator of Arabinose/Xylose Inducible Regulon in Sulfolobus acidocaldarius and Its Application for Homologous Protein Expression. Front. Microbiol. 11, 1066. doi:10.3389/fmicb.2020.01066.

van Wolferen, M., Ajon, M., Driessen, A. J. M., and Albers, S.-V. (2013) Molecular analysis of the UV-inducible pili operon from Sulfolobus acidocaldarius. Microbiologyopen 2, 928–937. doi:10.1002/mbo3.128.

van Wolferen, M., Ma, X., and Albers, S.-V. (2015) DNA Processing Proteins Involved in the UV-Induced Stress Response of Sulfolobales. J. Bacteriol. 197, 2941–2951. doi:10.1128/JB.00344-15.

van Wolferen, M., Wagner, A., van der Does, C., and Albers, S.-V. (2016) The archaeal Ced system imports DNA. Proc. Natl. Acad. Sci. U. S. A. 113, 2496–2501. doi:10.1073/pnas.1513740113.

Vassart, A., Van Wolferen, M., Orell, A., Hong, Y., Peeters, E., Albers, S.-V., et al. (2013) Sa-Lrp from Sulfolobus acidocaldarius is a versatile, glutamine-responsive, and architectural transcriptional regulator. Microbiologyopen 2, 75–93. doi:10.1002/mbo3.58.

Wagner, M., Shen, L., Albersmeier, A.Kolk, N. van der, Kim, S., Cha, J., et al. (2018) Sulfolobus acidocaldarius Transports Pentoses via a Carbohydrate Uptake Transporter 2 (CUT2)-Type ABC Transporter and Metabolizes Them through the Aldolase-Independent Weimberg Pathway. Appl. Environ. Microbiol. 84. doi:10.1128/AEM.01273-17.

Wagner, M., van Wolferen, M., Wagner, A., Lassak, K., Meyer, B. H., Reimann, J., et al. (2012) Versatile Genetic Tool Box for the Crenarchaeote Sulfolobus acidocaldarius. Front. Microbiol 3, 214. doi:10.3389/fmicb.2012.00214.

Wang, K., Sybers, D., Maklad, H. R., Lemmens, L., Lewyllie, C., Zhou, X., et al. (2019) A TetR-family transcription factor regulates fatty acid metabolism in the archaeal model organism Sulfolobus acidocaldarius. Nat. Commun. 10, 1542. doi:10.1038/s41467-019-09479-1.

Wang, L., Wang, S., and Li, W. (2012) RSeQC: quality control of RNA-seq experiments. Bioinformatics 28, 2184–2185. doi:10.1093/bioinformatics/bts356.

Ye, X., van der Does, C., and Albers, S.-V. (2020) SaUspA, the Universal Stress Protein of Sulfolobus acidocaldarius Stimulates the Activity of the PP2A Phosphatase and Is Involved in Growth at High Salinity. Front. Microbiol. 11. doi:10.3389/fmicb.2020.598821.

Zabriskie, T. M., and Jackson, M. D. (2000) Lysine biosynthesis and metabolism in fungi. Nat. Prod. Rep. 17, 85–97. doi:10.1039/a801345d.

