## Supplementary Information (Main) for "Machine learning uncovers a data-driven transcriptional regulatory network for the Crenarchaeal thermoacidophile *Sulfolobus acidocaldarius*"

### Saci Manuscript Supplementary

Main Manuscript: Saci Manuscript Draft

#### Supplementary Figure 1

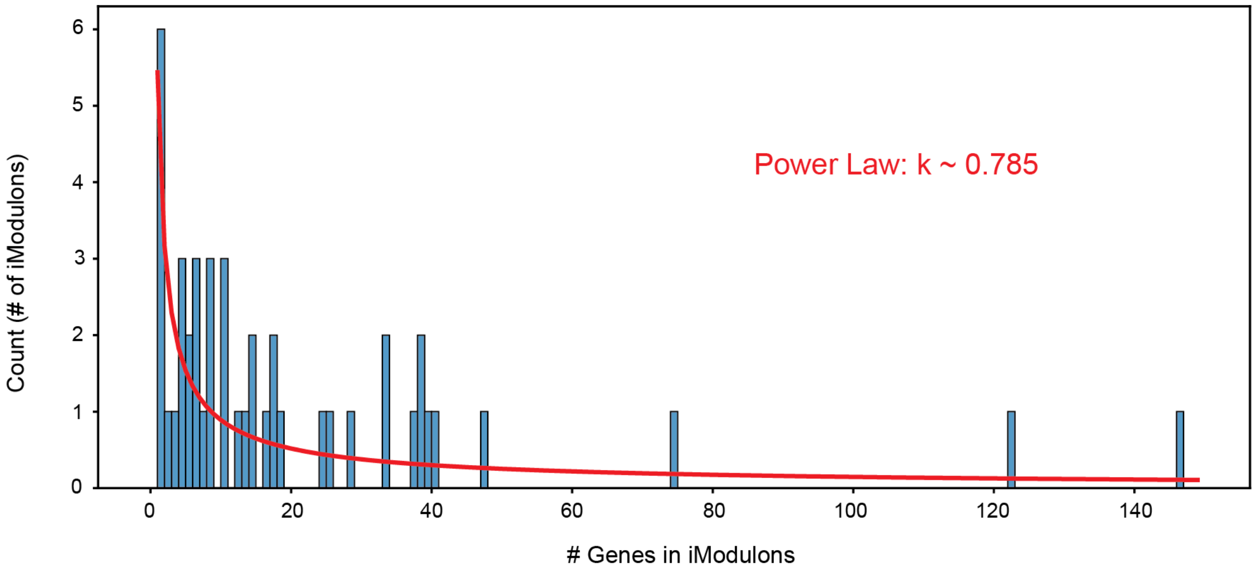


**Supplementary Figure 1:** Histogram of iModulons by gene set size (bin size of 1). Most iModulons contain below 20 genes, and all but 3 iModulons contain below 50 genes. Power Law formulation (with exponent parameter of ~0.785) overlaid this distribution in red.

### Supplementary Figure 2

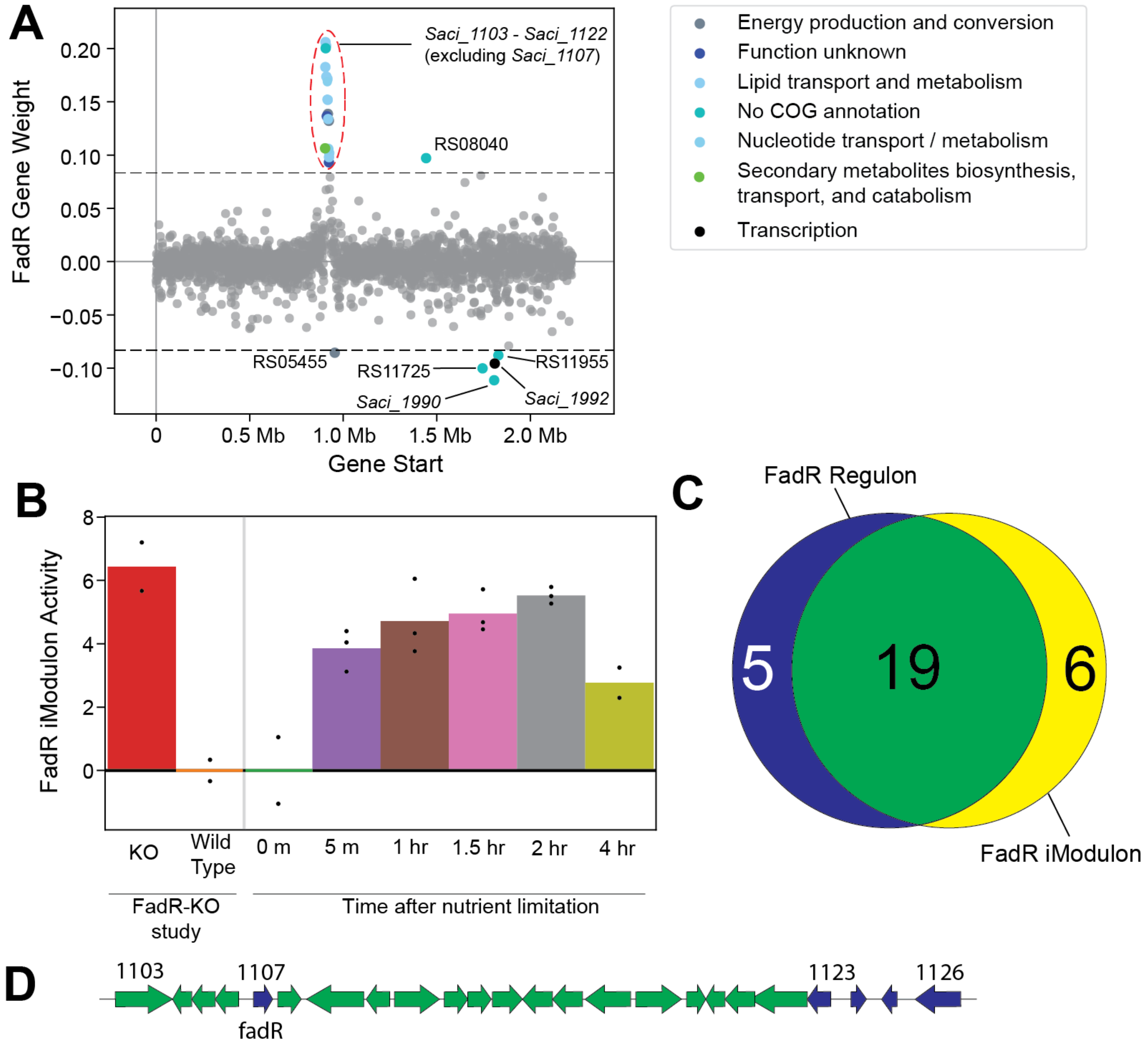


**Supplementary Figure 2:** Overview of the FadR iModulon. **a)** Scatter plots of the FadR iModulon gene weights. Genes are colored by COG categories. **b)** Barplot of FadR iModulon activity in 2 projects. **c)** Venn diagram comparing iModulon Recall with Regulon Recall. **d)** Gene map of FadR gene cluster in *S. acidocaldarius*. Green genes are in both iModulon and regulon, and blue genes are only in the regulon.

**Supplementary File 1:** A draft TRN (tsv format) assembled from extensive literature review. Each gene-regulation interaction is mapped towards a PMID listing the reference where it was first reported.

**Supplementary File 2:** A curated genome annotation (csv format) file

**Supplementary File 3:** InterPro scan results (json format) for uncharacterized genes in UV-tfb3 iModulon (*Saci_0951, Saci_1225, Saci_1270, Saci_1302,* and SACI_RS06010)

**Supplementary File 4:** InterPro scan results (json format) for putative peptide-induced D-xylose transporters (*Saci_0324*, *Saci_0675*, and *Saci_2095*)
